# A screen for combination therapies in *BRAF/NRAS* wild type melanoma identifies nilotinib plus MEK inhibitor as a synergistic combination

**DOI:** 10.1101/195354

**Authors:** Marco Ranzani, Kristel Kemper, Magali Michaut, Oscar Krijgsman, Nanne Aben, Vivek Iyer, Kim Wong, Theodoros I. Roumeliotis, Martin Del Castillo Velasco-Herrera, Jérémie Nsengimana, Gemma Turner, Nicola Thompson, Aida Shahrabi, Marcela Sjoberg, Mamunur Rashid, Anneliese O. Speak, Vera Grinkevich, Fiona Behan, David Tamborero, Francesco Iorio, Stijn van Dongen, Graham R. Bignell, Clara Alsinet, Sofia Chen, Emmanuelle Supper, Ken Dutton-Regester, Antonia Pritchard, Chi Wong, Anton Enright, Julia Newton-Bishop, Ultan McDermott, Nicholas K. Hayward, Jyoti S. Choudhary, Kosuke Yusa, Lodewyk Wessels, Mathew J. Garnett, Daniel Peeper, David J. Adams

**Affiliations:** Experimental Cancer Genetics, The Wellcome Trust Sanger Institute, Hinxton, Cambridge, CB10 1SA, United Kingdom; Division of Molecular Oncology, The Netherlands Cancer Institute, 1066 CX Amsterdam, the Netherlands; Division of Molecular Carcinogenesis, The Netherlands Cancer Institute, 1066 CX Amsterdam, the Netherlands; Mass Spectrometry, The Wellcome Trust Sanger Institute, Hinxton, Cambridge, CB10 1SA, United Kingdom, and Functional Proteomics Group, Chester Beatty Laboratories, The Institute of CancerResearch, London SW3 6JB, United Kingdom; Leeds Institute of Cancer and Pathology, University of Leeds, Cancer Genetics Building,St James's University Hospital, Beckett Street, Leeds, LS9 7TF, United Kingdom; Cancer Genome Project, The Wellcome Trust Sanger Institute, Hinxton, Cambridge, CB10 1SA, United Kingdom; Research Program on Biomedical Informatics, IMIM Hospital del Mar Medical Research Institute and Universitat Pompeu Fabra, Doctor Aiguader 88, 08003, Barcelona, Catalonia, Spain; European Molecular Biology Laboratory - European Bioinformatics Institute, Wellcome Genome Campus, Cambridge, CB10 1SD, United Kingdom; Oncogenomics Laboratory, QIMR Berghofer, Medical Research Institute, Brisbane, Queensland, Australia; Cellular Genomics, The Wellcome Trust Sanger Institute, Hinxton, Cambridge CB10 1SA, United Kingdom

## Abstract

Despite recent therapeutic advances in the management of *BRAF*^*V600*^-mutant melanoma, there is still a compelling need for more effective treatments for patients who developed *BRAF*/*NRAS* wild type disease. Since the activity of single targeted agents is limited by innate and acquired resistance, we performed a high-throughput drug screen using 180 drug combinations to generate over 18,000 viability curves, with the aim of identifying agents that synergise to kill *BRAF*/*NRAS* wild type melanoma cells. From this screen we observed strong synergy between the tyrosine kinase inhibitor nilotinib and MEK inhibitors and validated this combination in an independent cell line collection. We found that AXL expression was associated with synergy to the nilotinib/MEK inhibitor combination, and that both drugs work in concert to suppress pERK. This finding was supported by genome-wide CRISPR screening which revealed that resistance mechanisms converge on regulators of the MAPK pathway. Finally, we validated the synergy of nilotinib/trametinib combination *in vivo* using patient-derived xenografts. Our results indicate that a nilotinib/MEK inhibitor combination may represent an effective therapy in *BRAF*/*NRAS* wild type melanoma patients.

## INTRODUCTION

The development of targeted therapies has dramatically improved the treatment of *BRAF*^*V600E*^-mutant melanoma, with the combination of a BRAF and a MEK inhibitor achieving response rates of up to 68% and median progression-free survival of up to 12 months^1, 2^. On the contrary, for *BRAF*/*NRAS* wild type (WT) melanoma, which represents 25-30% of all melanoma cases^3, 4^, little progress has been made and there are currently no available targeted therapies. More recently, immunotherapies have revolutionized the treatment of melanoma with immune checkpoint inhibitors achieving response rates >60% and unprecedented long-term durable disease control, regardless of *BRAF* mutation status^5, 6,7^. Notwithstanding these advances, most melanoma patients are not cured by available therapies and second line treatments are required for patients who relapsed following immunotherapies.

High-throughput drug screens of cancer cell lines represent an effective approach to identify candidate compounds with high activity in specific subtypes of human cancer^8, 9^. Given the compelling need for new targeted therapies for patients with *BRAF*/*NRAS* WT melanoma, and the lack of large numbers of deeply characterized cell line models for this subtype of the disease, we assembled a large collection of *BRAF*/*NRAS* WT melanoma cell lines and characterised them for mutations, copy number alterations, and for their gene and microRNA expression profiles. We then performed a high-throughput drug combination screen in theselines generating over 18,000 viability curves with 180 drug combinations. We subsequently investigated the mechanism of synergy of the lead combination and used genome-wide CRISPR screening to pre-emptively identify mechanisms of drug resistance. Finally, we validated the efficacy of the combination *in vivo* using patient-derived xenografts.

## RESULTS

### Genomic and transcriptomic analysis of *BRAF*/*NRAS* wild type melanoma cell lines

We assembled a collection of 22 human melanoma cell lines, including 20 *BRAF*/*NRAS* WT, one *BRAF*^V600E^-mutant and one *NRAS*^Q61R^-mutant cell line (**Supplementary Table 1**) for screening. We catalogued somatic single nucleotide variants (SNV) in these lines by exome sequencing (**Supplementary Table 2**), copy number variation by use of SNP6 arrays (**Supplementary Table 3**), and gene and microRNA expression by RNA sequencing (**Supplementary Table 4-5-6**) (outlined in **Fig. 1a**). In agreement with the analysis of *BRAF*/*NRAS* WT human tumors from The Cancer Genome Atlas (TCGA) collection^4^, the *BRAF*/*NRAS* WT melanoma cell lines had a high mutational load (median 59.01 SNV/Mb, range 1.34-512.96; **Supplementary Fig. 1a** and **Supplementary Table 2b**) dominated by C>T mutations at dipyrimidines (**Supplementary Fig. 1c-d**, **Supplementary Table 2c**). *NF1* mutant cell lines displayed a significantly higher mutation frequency than cell lines without mutations in *BRAF, NRAS* or *NF1* (*P*<0.0001; One way Anova and Tukey’s multiple comparison test), recapitulating the pattern described in tumors from the TCGA and Yale Melanoma Genome Projects^3, 4^ (**Supplementary Fig. 1b, Supplementary Table 2b**). To identify putative melanoma driver genes we ran IntOGen^10^ using SNV data from 74 TCGA *BRAF*/*NRAS* WT tumors^4^ and found 24 genes that were statistically significantly mutated (**Supplementary Table 7a-b,** see **Methods**). Similarly, we collated melanoma drivers in regions defined as recurrently amplified or deleted in 333 melanomas from the TCGA collection^4^ spanning all cutaneous melanoma subtypes (**Supplementary Table 7c**). All 24 point mutated *BRAF*/*NRAS* WT melanoma drivers and 32 out of 39 driver genes in amplified or deleted regions were captured by somatic mutations/genomic alterations in at least one cell line in our collection (**Supplementary Fig. 1e, Supplementary Table 3b**). The mutation frequency of the 24 mutation drivers in our *BRAF*/*NRAS* WT cell line collection correlated with the frequency found in the aforementioned TCGA *BRAF*/*NRAS* WT tumors (Pearson correlation *P*<0.0001, R^2^ = 0.6198; **Fig 1b** and **Supplementary Fig. 1f**). Collectively, these data show that our cell line collection is representative of the major driver mutations found in *BRAF*/*NRAS* WT melanoma.

**Figure 1.**
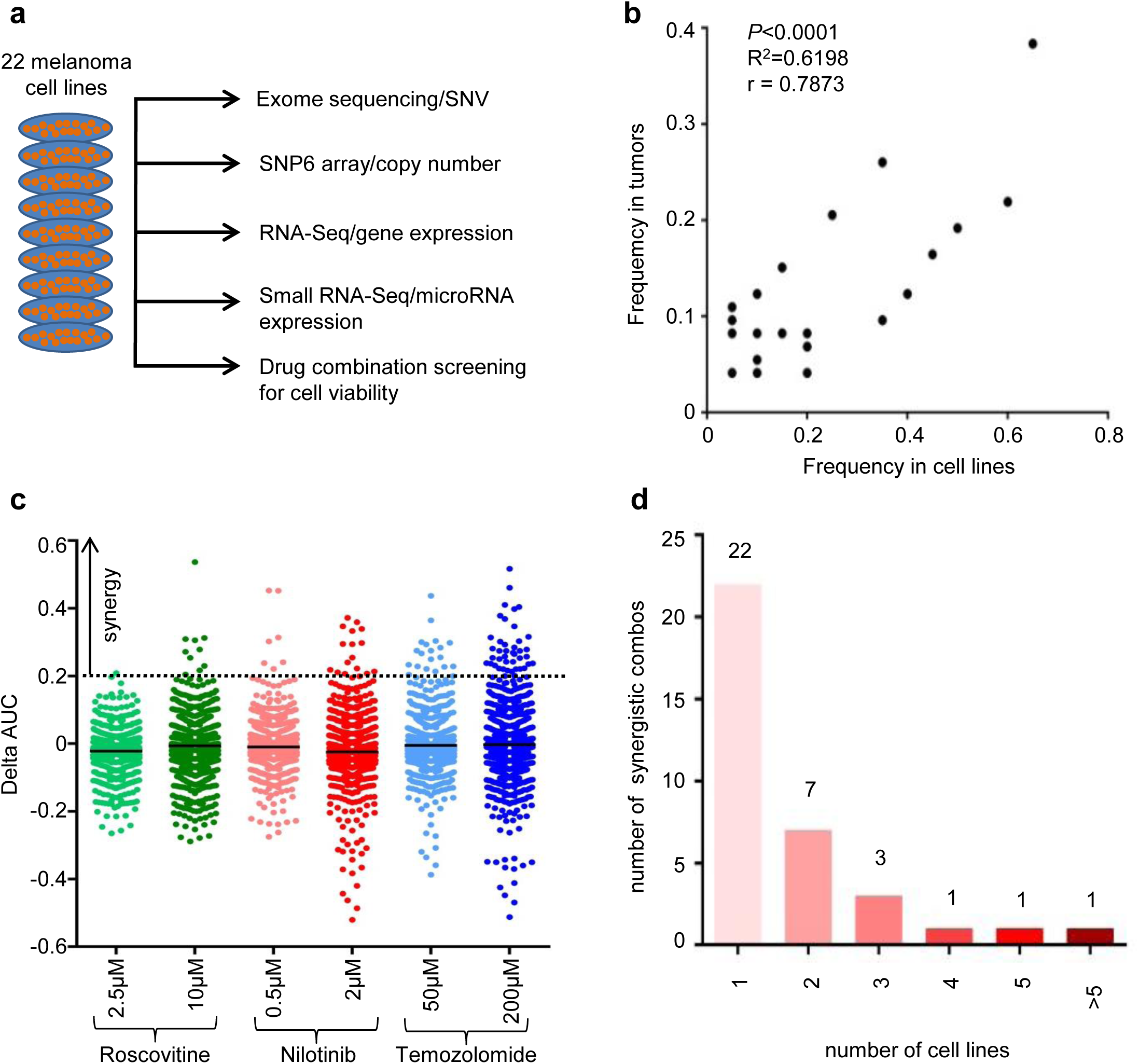
High-throughput drug screening of melanoma cell lines. **a)** Outline of data generated for the collection of 22 melanoma cell lines (see **Methods**). **b)** Correlation between the frequency of mutation in *BRAF*/*NRAS* WT melanoma drivers in the 20 *BRAF*/*NRAS* WT cell lines of our collection (X axis) and in 74 *BRAF*/*NRAS* WT tumors from the TCGA cohort^4^ (Y axis). *P* value, R^2^ and r by Pearson correlation. **c)** Delta AUC (Y axis, see **Methods**) of the library drugs combined with the different concentrations of the anchor drugs (X axis). Each dot represents the delta AUC of a drug combination in a cell line; the black line shows the mean, the dashed line the 0.2 delta AUC value. **d)** Number of combination that achieved a delta AUC>0.2 (Y axis) in recurrent (X axis) cell lines.See also **Supplementary Fig. 1** and **Supplementary Tables 1-9**.

### Nilotinib synergizes with MEK inhibitors in *BRAF*/*NRAS* wild type melanoma cell lines

Since our collection of *BRAF*/*NRAS* WT cell lines represents the largest of its kind we elected to perform a high-throughput drug combination screen with 60 library drugs, and three anchor drugs [temozolomide (alkylating agent), nilotinib (tyrosine kinase inhibitor) and roscovitine (broad CDK inhibitor) (**Supplementary Table 8**)] to identify candidate drug combinations for clinical use. Anchors were selected because of their broad, yet distinct, modes of action thus allowing us to cover a wide pharmacological space. Anchor drugs were tested at two concentrations and combined with each of the 60 library drugs tested at five concentrations over a 256-fold concentration range. Cellular viability was measured six days after drug treatment and normalized against DMSO-treated controls. Overall, we tested 180 drug combinations and generated 18,810 survival curves (three curves per cell line per combination, **Supplementary Table 9**). Survival curves were analyzed using the Area Under the Curve (AUC) method as described previously^8^ (**Supplementary Fig. 1g**). The viability of cell lines treated with the single library drug alone was generally higher than cell lines treated with drug combinations (average ± SEM of AUC is 0.8684 ± 0.001775 and 0.7925 ± 0.002426 for library drugs and drug combinations, respectively; *P*<0.0001 by unpaired Student’s t-test) (**Supplementary Fig. 1h-j**). To prioritize the most effective drug combinations, we measured drug synergy as the difference between the AUC of the predicted additive effect^11^ and the AUC of the drug combination (delta AUC) (**Supplementary Fig. 1g**; see **Methods**). With a threshold of delta AUC>0.2 (selected to represent the top 1.5% of 6270 delta AUCs, see **Methods**), we identified 94 occurrences of synergy from 53 drug combinations (**Fig. 1c, Supplementary Table 9d**). Most combinations showed synergy in a single cell line; six combinations displayed synergy in 3 or more cell lines, 3.3% (6/180) of all combinations tested (**Fig. 1d** and **Supplementary Table 9e**).

We triaged these combinations as follows. Firstly, we assayed the synergy observed in the high-throughput screen in the three cell lines with the highest delta AUC using a low-throughput viability assay (see **Methods**). We focused on the five drug combinations with the highest average delta AUC where synergy was observed in three or more cell lines (**Fig. 1d** and **Supplementary Table 9e**). Dose response curves were performed in triplicates using the same 256-fold doses range used in the high-throughput screen. In this way, we confirmed synergy between temozolomide and olaparib (PARP inhibitor) and between nilotinib and PD-0325901 (MEK inhibitor) (**Fig. 2a, Supplementary Table 10a,** see **Methods** for delta AUC threshold). Testing of the three combinations just below the defined threshold (delta AUC>0.2 in two cell lines only, see **Methods**) did not support the results of the high-throughput screen (**Supplementary Table 10a**).

**Figure 2.**
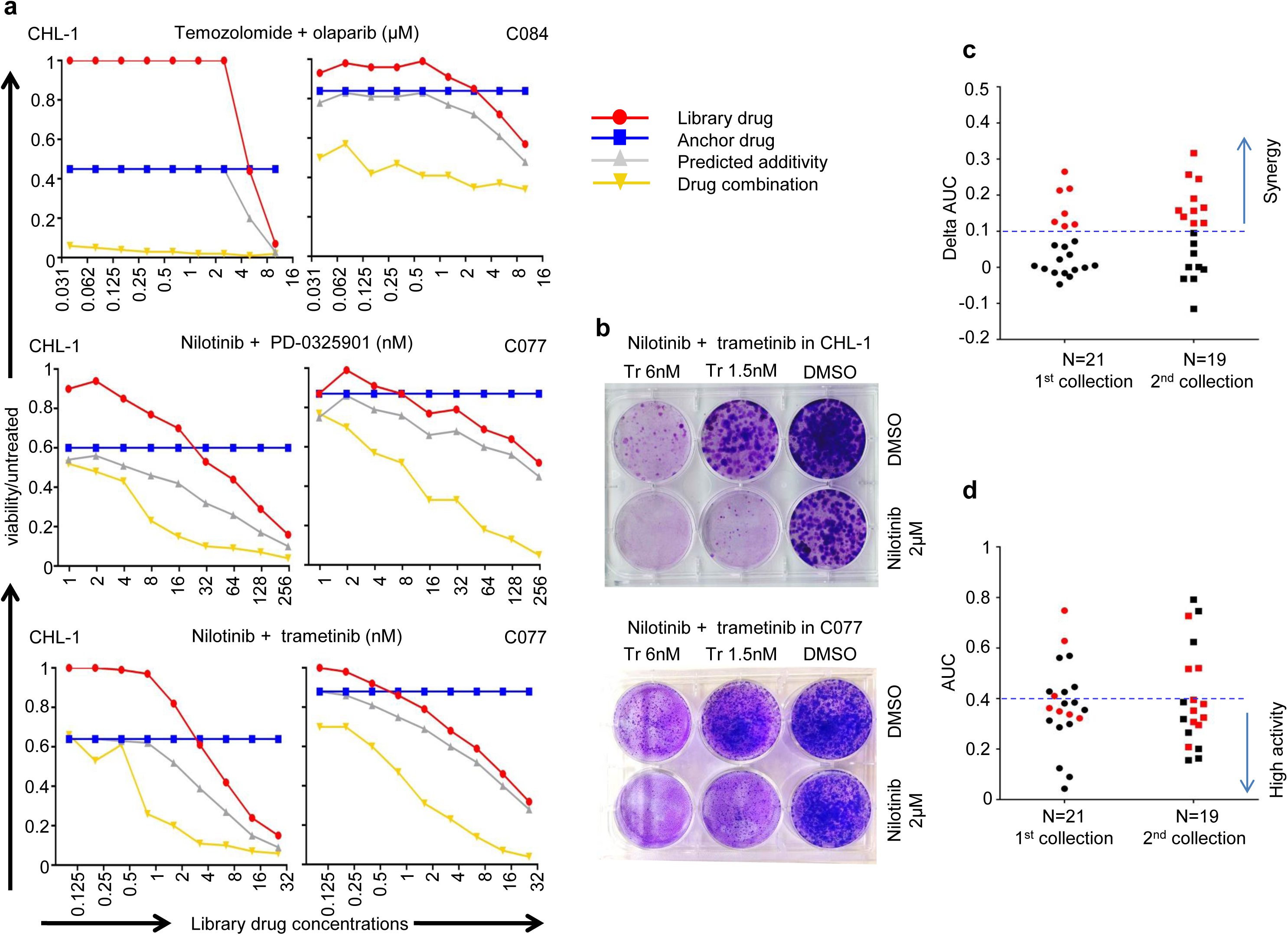
Confirmation of the synergy between nilotinib and MEK inhibitors in *BRAF*/*NRAS* WT melanoma. **a)** Survival curves of two representative cell lines treated with temozolomide plus olaparib (top panels, representative of two independent experiments), nilotinib plus PD-0325901 (middle panel, representative of three independent experiments) and nilotinib plus trametinib (bottom panel, representative of three independent experiments). The Y axis shows viability vs vehicle treated control, X axis the concentration (μM for olaparib, nM for PD-0325901 and trametinib) of the library drug. The red line shows the viability of the cells treated with the library drug alone, the blue line the viability with the anchor drug alone, the yellow line the viability with the drug combination, the grey line the predicted additivity (see **Methods**). Each point is the average value of a technical triplicate. **b)** Clonogenic assays confirming the synergy between nilotinib and MEK inhibitors. The concentrations of the library and anchor drugs are indicated on the top or on the right, respectively. These assays are representative of a biological duplicate. **c**-**d)** Summary of the delta AUC (c) and AUC combination (d) values for nilotinib/trametinib combination in the two collections of cell lines. The dashed lines represent values of delta AUC=0.1 and AUC=0.4 (see **Methods**). Red symbols indicate cell lines with delta AUC>0.1. See also **Supplementary Fig. 1-2** and **Supplementary Tables 10-11**.

We next repeated the low-throughput viability assays on 21 of the previously screened cell lines (including 19 *BRAF*/*NRAS* WT, one *BRAF*^V600E^-mutant and one *NRAS*^Q61R^-mutant cell lines, see **Supplementary Table 1**) using the drug combinations temozolomide with olaparib, nilotinib with PD-0325901 and nilotinib with trametinib (a second MEK inhibitor in clinical use^12^). The synergies between temozolomide and olaparib and between nilotinib and both MEK inhibitors were confirmed in 12 and 4 out of 19 *BRAF*/*NRAS* WT cell lines, respectively (**Fig. 2a**, **Supplementary Table 10b** and **Supplementary Table 11**, delta AUC>0.1, see **Methods** for synergy thresholds). Intriguingly, synergy was also observed in an *NRAS* mutant line analyzed in parallel. We further confirmed the potency of these drug combinations by performing clonogenic assays (**Fig. 2b** and **Supplementary Fig. 2a**).

Given the limited activity of alkylating agents combined with PARP inhibitors in clinical trials^13, 14^, we focussed on the nilotinib/trametinib combination for testing in a second independent collection of 19 melanoma cell lines, including 10 *BRAF*^V600^-mutant, 4 *NRAS*^Q61^-mutant and 5 *BRAF*/*NRAS* WT lines (**Supplementary Table 1b**). We observed synergy (delta AUC>0.1, see **Methods** for synergy threshold) in 10 out of 19 cell lines (**Supplementary Table 10c**). Synergies were confirmed in clonogenic assays (**Supplementary Fig. 2b**). Overall, we tested the nilotinib/trametinib combination in two collections of melanoma cell lines and found that it was synergistic in 17 out of 40 lines (42.5%, **Fig. 2c**), including 6/24 *BRAF*/*NRAS* WT cell lines. In 26 out of 40 (65%) melanoma cell lines we observed high activity of the drug combination (AUC<0.4, see **Methods**) as a result of synergy, additivity and single agent activity (**Fig. 2d**).

### AXL expression is associated with synergy between nilotinib and MEK inhibitors in *BRAF/NRAS* wild type melanoma

To identify markers that predict synergy between nilotinib and MEK inhibitors, we looked for an association between the drug synergy score (delta AUC), coding mutations, copy number alterations and/or gene/microRNA expression (see **Methods**). To reduce multiple testing, we considered only lesions that were previously characterized as cancer drivers (**Supplementary Table 12a**) following an approach described previously^8^. We classified each lesion as a gain-of-function or loss-of-function alteration partitioning them into functional groups (see **Methods**) but did not identify any statistically significant gene/drug associations in this way (**Supplementary Table 13a**). We then extended our analysis to all lesions in melanoma drivers (**Supplementary Table 12b**), but again no associations were found (**Supplementary Table 13b**). Analysis of differentially expressed microRNAs (**Supplementary Table 5b-c**, see **Methods)** also did not find significant associations (**Supplementary Table 13c**).

We then looked for pathways differentially expressed between cell lines that showed synergy (sensitive) or no synergy (non-sensitive) to nilotinib combined with both MEK inhibitors (see **Methods** for cell line classification into sensitive and non-sensitive and **Supplementary Table 10b**). The sensitive cell lines displayed higher levels of cell cycle genes and lower levels of genes associated with pigmentation (**Supplementary Fig. 3a-b** and **Supplementary Table 14a-c,** see **Methods)**. This gene expression pattern was observed in ∼30% of tumors from two melanoma cohorts and was more frequent in tumors classified as “Proliferative” by the Jonsson’s gene expression classifier^15^ (**Supplementary Fig. 3c-d**). We next looked for genes expressed (FPKM>1 by RNA Sequencing) exclusively in the sensitive cell lines and found 4 transcripts, among which *AXL* displayed the highest differential expression (314.28-fold change in sensitive vs non-sensitive; **Supplementary Table 14d**). We considered this finding an interesting connection, because we and others have previously shown that *AXL* is involved in resistance to targeted therapies in melanoma^16, 17^. Therefore, we further investigated its association with the response to nilotinib/trametinib treatment. We confirmed the previously described inverse correlation between *AXL* and *MITF* RNA expression^17^ in our *BRAF*/*NRAS* WT cell lines (*P*= 0.0013, R^2^ = 0.4842, by Pearson correlation; **Supplementary Fig. 3e** and **Supplementary Table 15a**) and found that *AXL* expression levels were significantly higher in sensitive cell lines compared to non-sensitive cell lines (*P*=0.0117 by unpaired Student’s t-test, **Supplementary Fig. 3f-g** and **Supplementary Table 15a**), as expected. We next measured protein expression in sensitive cell lines and representative non-sensitive cell lines revealing that all *BRAF*/*NRAS* WT sensitive cell lines in our collection expressed AXL, whereas it was undetectable in non-sensitive lines (**Fig. 3a, Supplementary Fig. 3h**). Extending the analysis to the *BRAF*/*NRAS* WT non-sensitive cell lines of the second collection, we found that 4/5 non-sensitive cell lines were AXL^neg^, with only 1/5 non-sensitive cell line expressing AXL protein (**Supplementary Fig. 3i**). Overall, AXL^pos^ cell lines were significantly enriched for the occurrence of synergy between nilotinib and trametinib (*P* = 0.007 by two tailed Fisher’s exact test, **Supplementary Table 15c**). Remarkably, although AXL expressing cell lines displayed higher synergy for the nilotinib/trametinib combination, they showed higher resistance to MEK inhibitors alone (**Supplementary Fig. 3j**), in agreement with previous studies^16, 17^. Notably, we did not observe a clear association between *AXL* expression and synergy in *BRAF*^V600^ or *NRAS*^Q61^-mutant cell lines (**Supplementary Table 15b** and **Supplementary Fig. 3h**).

**Figure 3.**
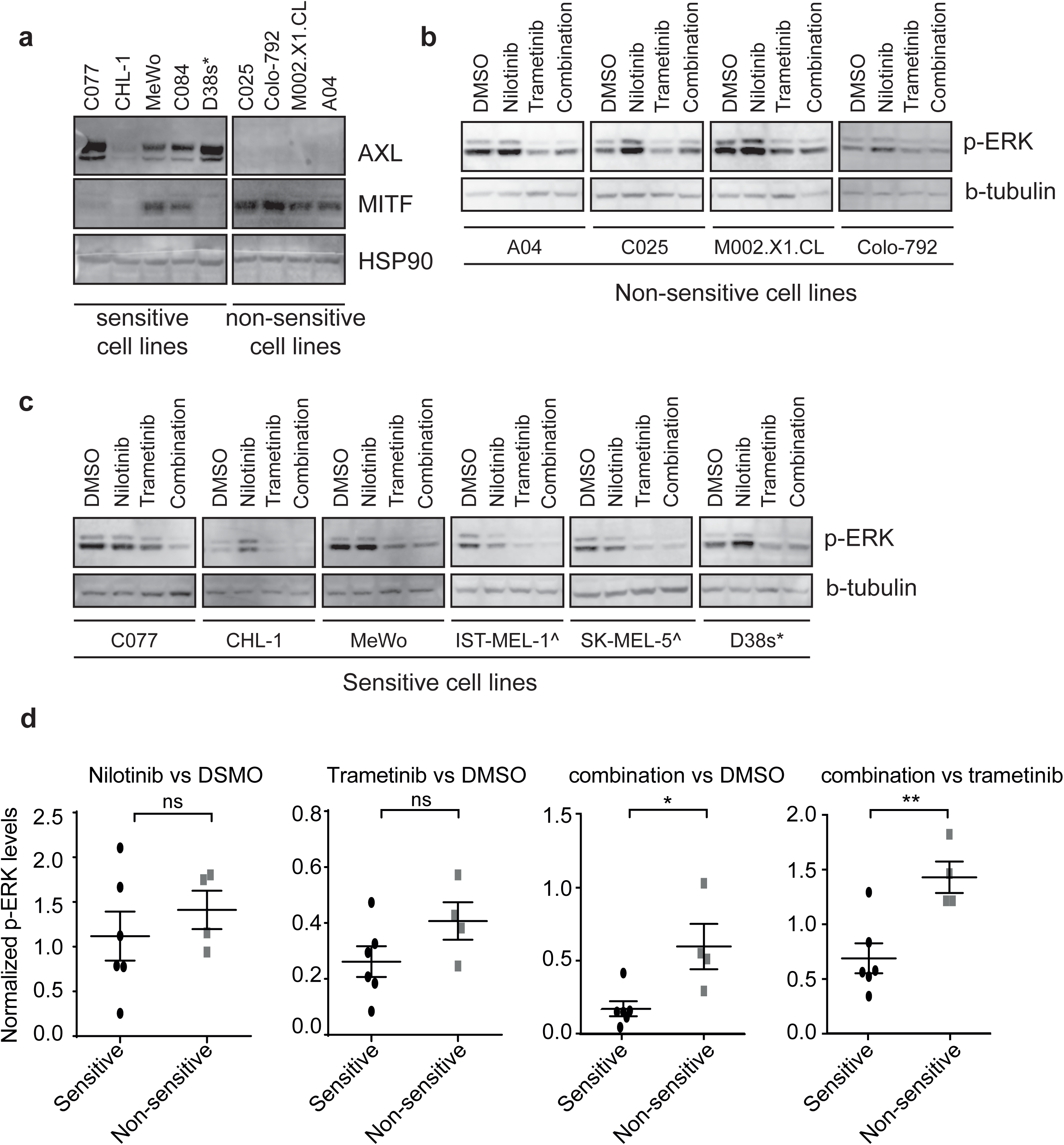
AXL expression and p-ERK downregulation are associated with the synergy between nilotinib and MEK inhibitors. **a)** Western blot (WB) for AXL and MITF expression in 5 sensitive and 4 non-sensitive cell lines. HSP90 is displayed below as loading control. **b-c)** Western blot for p-ERK in non-sensitive (b) and sensitive (c) cell line upon treatment with DMSO vehicle, nilotinib (2μM), trametinib (1nM) or combination for 6h (see **Methods**). Beta tubulin (β-tubulin) loading control is displayed below the blot. The blot is representative of experiments conducted in biological triplicate. * indicates *NRAS*^*Q61R*^-mutant, ^ indicates *BRAF*^*V600E*^-mutant cell lines. **d)** Level of p-ERK quantified by WB in sensitive and non-sensitive cell lines. Each cell line was assayed in 3 independent experiments. For each experiment, the levels of p-ERK were normalized for the loading control, then the average per line was calculated among the 3 experiments (each point in the dot plots). The dot plot shows the average and the standard error mean of the group (sensitive/non-sensitive cell lines). Significance is calculated by unpaired Student’s t-test. The analysis of the images was performed by an operator blinded to the sample identifiers. See also **Supplementary Fig. 3** and **Supplementary Tables 12-15**.

We then investigated the effect of the perturbation of AXL expression on nilotinib/trametinib synergy in *BRAF*/*NRAS* WT melanoma cell lines. We first confirmed that infection with lentiviral vector expressing *AXL* cDNA or short hairpin RNA targeting *AXL* induced up and downregulation of AXL protein, respectively (**Supplementary Fig. 3k**). The overexpression of AXL did not affect nilotinib/trametinib synergy in non-sensitive cell lines (**Supplementary Fig. 3l-m** and **Supplementary Table 16**). Conversely we found that AXL knockdown reduced, but did not fully abrogate, the effect of nilotinib/trametinib in sensitive cell lines (*P* = 0.084 and *P* = 0.025 in C077 and MeWo cell lines, respectively **Supplementary Fig. 3l-m** and **Supplementary Table 16**). This result is in keeping with the view that AXL expression is associated with a transcriptional cell state involved in drug resistance rather than being the only functional regulator of resistance *per se*^16, 17^.

### The synergy between nilotinib and trametinib is associated with increased pERK repression

To further dissect the mechanism of the observed synergy, we performed a high-throughput proteome and phosphoproteome analysis of representative sensitive (C077) and non-sensitive (C025) *BRAF*/*NRAS* WT cell lines treated with trametinib, nilotinib or the nilotinib/trametinib combination (**Supplementary Fig. 3n**, see **Methods**). These data were used to assess the effects of drug treatment on the protein phosphorylation landscape. Among the 9657 detected phospho-peptides (see **Methods**), we found 1753 phospho-sites that were significantly altered with either trametinib, nilotinib or the trametinib/nilotinib combination (**Supplementary Table 17a**). Intriguingly nilotinib did not decrease the phosphorylation of any of its known targets^18^, but instead induced the upregulation of pRAF1 and pERK1 (**Supplementary Table 17b-c**). Indeed, a closer analysis of pERK1/2 levels showed that nilotinib induced pERK in both cell lines (**Supplementary Fig. 3o** and **Supplementary Table 17b-c)**, a phenomenon potentially explained by its activity as a mild RAF inhibitor that stimulates RAFs and MEKs activity inducing paradoxical ERK activation^19, 20^. Conversely, the combination of trametinib and nilotinib induced repression of pERK in the sensitive cell line but not in the non-sensitive cell line (**Supplementary Fig. 3o** and **Supplementary Table 17b-c).** These results from phosphoproteomics were validated in 6 sensitive and 4 non-sensitive cell lines by Western blotting where sensitive cell lines treated with the combination displayed a reduction of pERK that was significantly more pronounced when compared tonon-sensitive cell lines (*P*=0.0154 and *P*=0.0069 vs vehicle-treated or trametinib-treated, respectively; unpaired Student’s t-test) (**Fig. 3b-d** and **Supplementary Fig. 3p**). Total ERK did not change (**Supplementary Fig. 3q**). Our results showed that trametinib and nilotinib work synergistically to silence pERK and suppress melanoma cell growth.

### Resistance to nilotinib/trametinib occurs via regulators of MAPK signalling

While combinatorial treatment allows for more durable clinical responses to therapy^1, 2^, eventually the emergence of resistance can cause cancer relapse in most patients^21^. Therefore, we decided to investigate the molecular mechanisms that might mediate resistance to the nilotinib/trametinib combination by use of CRISPR/Cas9 screens. We generated three Cas9 expressing cell lines (CHL-1, C077 and MeWo), which were selected because they displayed sensitivity to the nilotinib and trametinib combination, and transduced them with a genome-wide sgRNA library^22^. Cells were cultured for 18 days in trametinib, nilotinib/trametinib, or DMSO vehicle (**Supplementary Fig. 4a-d**, see **Methods**). We observed only limited overlap of genes whose loss conferred resistance to the nilotinib/trametinib combination among the three cell lines (**Fig. 4a**, **Supplementary Fig. 4e** and **Supplementary Table 18-20,** see **Methods**). We also found limited overlap among genes conferring resistance to trametinib (**Supplementary Fig. 4f-g, Supplementary Tables 19-20**). This suggests that many different genes are potentially operative in mediating resistance and interact with the different genetic backgrounds of each cell line.

**Figure 4.**
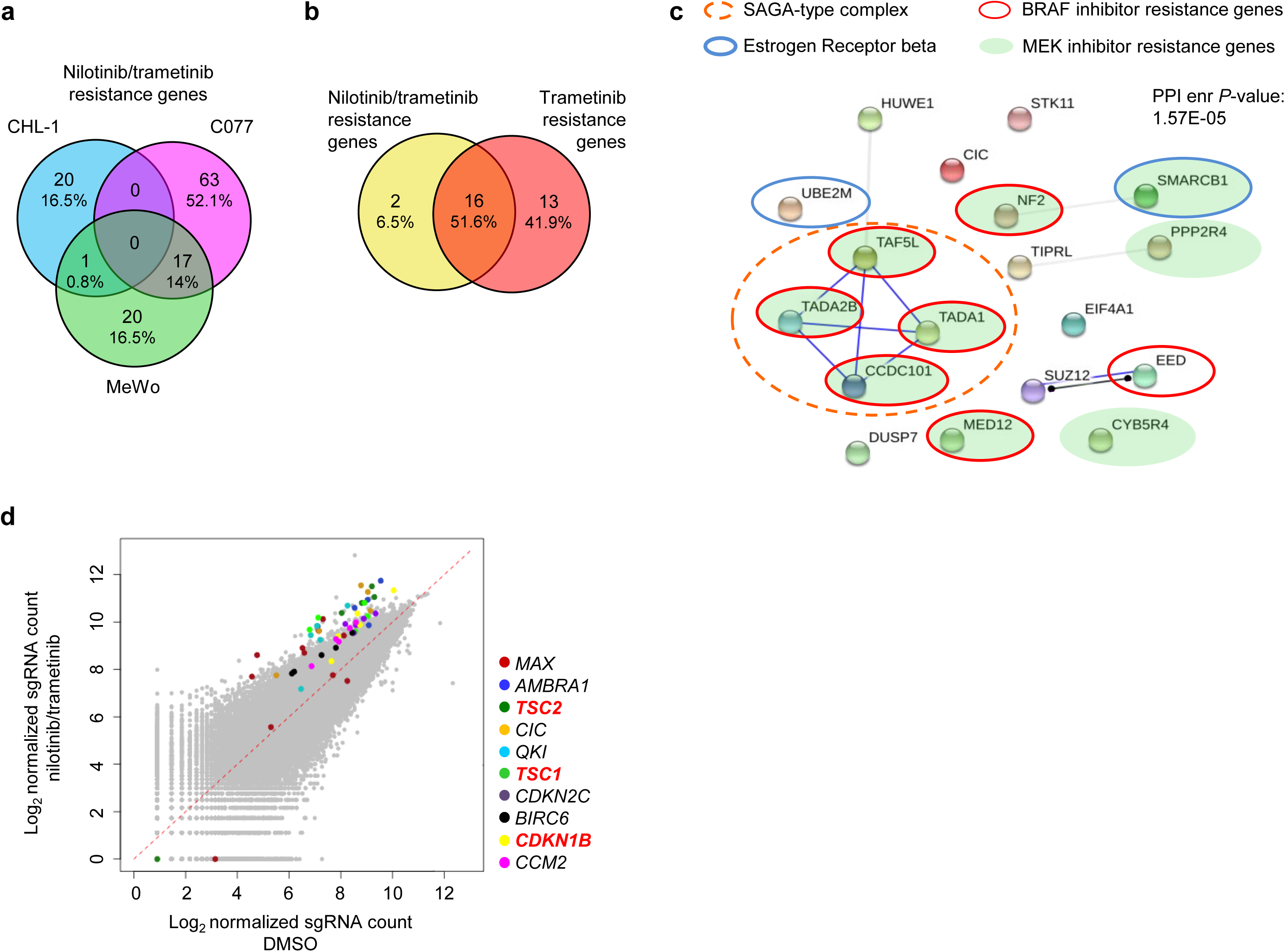
Identification of the mechanisms of drug resistance to the nilotinib/trametinib combination by CRISPR-Cas9 genome-wide library screening. **a)** Venn diagram of genes conferring resistance to the drug combination in CHL-1, C077, MeWo cell lines; number of genes and % of total are indicated. We considered genes with FDR<0.1 (by MAGeCK, see **Methods**) in both replicates. **b)** Venn diagram of the genes conferring resistance to the nilotinib/trametinib combination or trametinib in 2 or more cell lines (criteria as in a). **c)** Network of protein-protein interaction for the genes conferring resistance to nilotinib/trametinib combination (criteria as in b). Blue lines indicate binding, black lines reaction, grey lines unspecified interaction; PPI enr is the protein-protein interaction enrichment *P-*value calculated by STRING (see **Methods**). The coloured circles highlight genes belonging to top enriched pathways (see **Supplementary Table 19a**). **d)** Log_2_ of the normalized sgRNA count (see **Methods**) for each gRNA in vehicle treated (X axis) and drug combination treated (Y axis) CHL-1 cells after 18 days of treatment. The different sgRNAs targeting each of the top 10 enriched genes are color coded as detailed in the legend. See also **Supplementary Fig. 4** and **Supplementary Tables 18-21**.

Given the heterogeneity of the observed mechanisms of resistance, we focussed on the genes that conferred drug resistance in at least two of the three cell lines. Interestingly, we observed that many combination resistance genes interact (*P*=1.57 10^−9^ by STRING^23^) and that these genes are significantly enriched for members of the SAGA-type complex, estrogen receptor beta network and chromatin regulators (**Fig. 4c**, **Supplementary Table 21a**). In line with the converging inhibitory activity on the MAPK-ERK axis, we observed a large overlap between nilotinib/trametinib resistance genes and trametinib-only resistance genes (**Fig. 4b**). Accordingly, 7/18 nilotinib/trametinib resistance genes have previously been identified as vemurafenib (BRAF inhibitor) resistance genes, while 9/18 genes have previously been identified as selumetinib (MEK inhibitor) resistance genes^24, 25^ (**Fig. 4c, Supplementary Fig. 4h** and **Supplementary Table 18k**). Notably, fewer genes (18 vs 29) appeared to confer resistance to the combination compared to trametinib alone (**Fig. 4b**), suggesting that the combination may overcome some mechanisms of resistance observed with trametinib alone.

### Nilotinib and trametinib synergise *in vivo* in two *BRAF/NRAS* wild type melanoma models

We next tested the nilotinib/trametinib combination *in vivo*. Firstly, we inoculated NOD.Cg-*Prkdc*^scid^ *Il2rg*^tm1Wjl^/SzJ (NSG) mice with the sensitive cell line MeWo and after tumor establishment we treated these mice with vehicle, trametinib, nilotinib or the nilotinib/trametinib combination by gavage (n=5 mice, n=10 tumors per each of the four treatment groups, see **Methods**). The combination induced a significant reduction of tumor growth compared to vehicle, nilotinib only and trametinib only treatments (*P* value = 0.0004, 0.0005, 0.004, respectively by unpaired Student’s t-test at the last time point, **Fig. 5a**). To validate the drug combination in a model that more closely represents human tumors^26, 27^, we interrogated a collection of *BRAF*/*NRAS* WT melanoma patient derived xenografts (PDX)^28^ for expression of AXL and MITF (**Fig. 5b**). Since AXL^pos^ cell lines are enriched for synergy between nilotinib and trametinib (**Fig. 3a** and **Supplementary Table 15c**), we selected the PDX line M003 which expressed the highest level of *AXL* for *in vivo* studies (**Fig. 5-c**). In the *in vivo* experiments (performed as described above for MeWo), nilotinib induced a mild reduction of tumor growth, trametinib induced a more pronounced tumor growth reduction and the combination induced partial regression, with a reduction in tumor volume that was maintained for the duration of the experiment (39 days of treatment; *P*<0.0001 for combination treated mice vs the three other groups by unpaired Student’s t-test at the last common time point; **Fig. 5d**). Analyses of representative tumors (n=4) at the experimental endpoint confirmed that the drug combination induced a significant reduction of p-ERK (**Fig. 5e**), as previously observed in cell lines (**Fig. 3c-d**), and caused an alteration of the tumor cellular morphology (**Fig. 5f**). These results confirmed the synergy between nilotinib and trametinib in two *in vivo* models of *BRAF*/*NRAS* WT melanoma.

**Figure 5.**
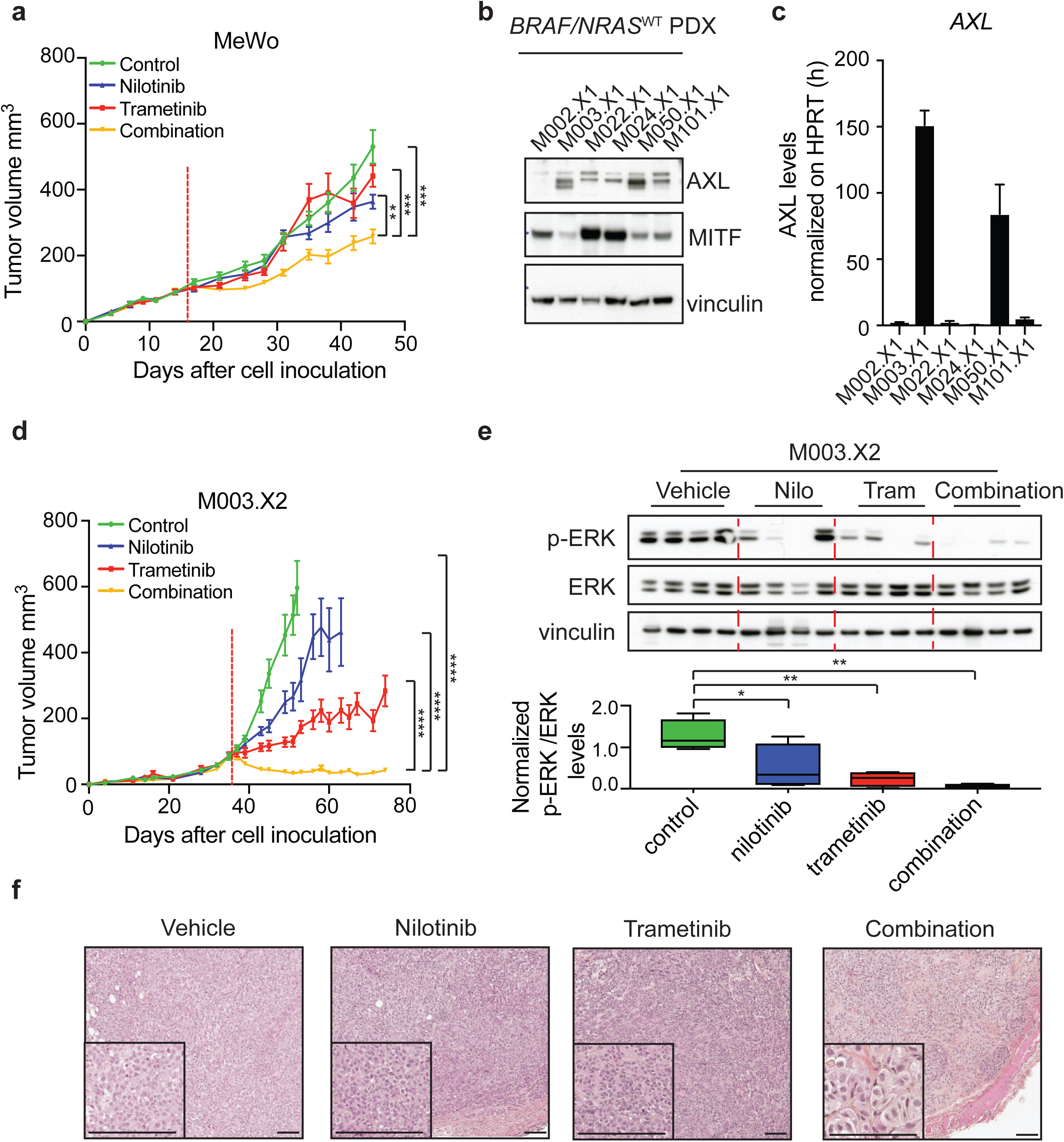
The combination of nilotinib plus trametinib is synergistic in two *in vivo* models. **a)** Volume (Y axis) of tumors from MeWo cell line inoculated in NOD.Cg-*Prkdc*^scid^ *Il2rg*^tm1Wjl^/SzJ (NSG) mice upon treatment with vehicle (green), nilotinib 75mg/kg/day (blue), trametinib 0.1mg/kg/day (red) or their combination (yellow) (n=10 tumors/group, see **Methods**). The graph shows the mean and the standard error mean. The vertical dashed red line highlights the start of the treatment. *P*-value calculated by unpaired Student’s t-test on the last time point; *\*P*<0.05*, **P*<0.01, \*\*\*\**P*<0.0001. **b**) Western blot for AXL, MITF and vinculin loading control in a collection of *BRAF*/*NRAS* WT melanoma PDX. **c)** Quantification of *AXL* RNA expression by Q-PCR in *BRAF*/*NRAS* WT PDX. **d**) Volume (Y axis) of tumors from M003.X2 PDX inoculated in NSG mice upon treatment with vehicle (green), nilotinib 75mg/kg/day (blue), trametinib 0.3mg/kg/day (red) or their combination (yellow). Graph as in a). *P*-value calculated by unpaired Student’s t-test at 51 days, the last time point when all the 4 experimental cohorts were viable; \*\*\*\**P*<0.0001. The suffix .X1-.X2 indicates the passage number of the PDX line. **e)** Top panel: western blot for p-ERK, total ERK and vinculin loading control in 4 representative M003.X2 tumors per group of treatment (indicated above the plots) collected at the experimental endpoint. Bottom panel: quantification in p-ERK levels (Y axis, normalized for total ERK) from the western blot displayed above. Box plot extends from the 25th to 75th percentiles, whiskers from min to max, the middle line indicates the median. *P*-value by one way Anova and Tukey’s multiple comparisons test; *\*P*<0.05*, **P*<0.01 **f)** Representative hematoxylin and eosin stained section of a tumor per each treatment group collected at the experimental endpoint. Scale bar=100μm, the bottom left corner displays a higher magnification.

## DISCUSSION

We assembled a collection of *BRAF*/*NRAS* WT melanoma cell lines and deeply characterized them to catalogue SNVs, CNVs, gene and microRNA expression, and their growth in response to 180 drug combinations. We show that our collection is representative of the mutations carried by *BRAF*/*NRAS* WT melanoma tumors (**Figure 1b** and **Supplementary Fig. 1a-f**) and thus represents a publicly available resource for hypothesis testing and model choice for *BRAF*/*NRAS* WT melanoma, which has been less widely studied when compared to *BRAF-* and *NRAS-* mutant disease. Analysis of the aforementioned high-throughput drug screen data revealed that synergy between anti-cancer drugs is rare, and in many cases private to a specific cell line. By applying a stringent and multi-step validation approach, we confirmed robust synergistic interactions between temozolomide and olaparib, and also a combination of nilotinib and MEK inhibitors. Since the combination of alkylating agents and PARP inhibitors has failed to elicit clinical benefit in melanoma clinical trials^13, 14, 29^, we focussed our efforts on the nilotinib/trametinib combination. Notably, nilotinib and trametinib are approved for the treatment of leukemia and melanoma, respectively^12^, and we detected synergy at concentrations far below the peak of plasma concentration achieved in patients^12,30, 31^. Additionally, our results in mouse models show that the nilotinib/trametinib combination can be tolerated *in vivo* with a regimen that induced regression in a PDX model of *BRAF/NRAS* WT human melanoma.

In sum, the nilotinib/trametinib combination showed synergy (AUC>0.1) in 42.5% (17/40) of all melanoma cell lines including 6/24 *BRAF/NRAS* wild type lines. Further, we also observed strong activity of the drug combination (AUC <0.4) in 65% (26 out of 40) of lines, including 62.5% (15 out of 24) of our *BRAF*/*NRAS* WT lines. Collectively, this effect on melanoma cell growth is the result of high single agent activity, additivity, and synergy and suggests that the combination could benefit a broad range of patients.

Notably, we did not identify a biomarker linked to drug synergy from the genomic data but discovered that AXL expression was associated with synergy between nilotinib and trametinib in *BRAF*/*NRAS* WT cell lines. AXL expression was frequently found in *BRAF*/*NRAS* WT PDX (five out of six, **Fig. 5b**) and is reported in a significant fraction of melanomas^17, 32^, thus suggesting that a sizeable fraction of patients with *BRAF*/*NRAS* WT melanoma may benefit from the nilotinib/trametinib combination. Previous studies have suggested that AXL expression is associated with a phenotype switch of melanoma cells towards a transcriptional status associated with drug resistance^16, 17^. In agreement, we observed that *BRAF*/*NRAS* WT AXL^pos^ cell lines display resistance to MEK inhibitors alone, yet display sensitivity to the nilotinib/trametinib combination. Interestingly, despite the overexpression or knockdown of AXL was not sufficient to revert the sensitive/non-sensitive status, in agreement with previous findings^16^, we observed that AXL knockdown reduced the synergy between nilotinib and trametinib. Overall, our and previous studies suggest a complex involvement of AXL in drug sensitivity, with AXL basal expression indicative of a cancer transcriptional status associated with differential drug sensitivity, rather than the exclusive determinant of the phenotype itself^16, 17^.

Given the promising results obtained by combining targeted and immune therapy^33, 34, 35^, we also envision the possible use of the nilotinib/trametinib combination with immune checkpoint inhibitors to improve patient outcome and disease control, or as a second line treatments following relapse after immune checkpoint therapy. Notably, the nilotinib/trametinib combination did not induce PD-L1 in *BRAF*/*NRAS* WT melanomas (n=4), suggesting that the treatment does not provoke an immune-suppressive phenotype (**Supplementary Fig. 5**) and might be compatible with immune-checkpoint blockade.

We undertook an unbiased high-throughput phosphoproteome approach to shed light on potential mechanisms of synergy between nilotinib and trametinib. Mechanistically we revealed that the synergy is due to the potent reduction of ERK phosphorylation. Since our and previous findings indicate that nilotinib inhibits RAFs^19, 20^, and trametinib is a well-established MEK1/2 inhibitor, our results suggest that the 2 drugs synergise by blunting ERK activation. This potent pERK inhibition was confirmed *in vivo* in a PDX model and further supported by the CRISPR/Cas9 screening in sensitive lines which revealed that nilotinib/trametinib resistance can be mediated by the same genes responsible for resistance to other MAPK pathway inhibitors^24, 25^.

Among the common nilotinib/trametinib resistance genes, we found 7 previously identified vemurafenib resistance genes^24^, including members of the SAGA complexes, *MED12* and *NF2*, and also regulators of the estrogen beta pathway which have anti-proliferative activity in melanoma^36, 37^. Some of these genes are mutated in a fraction of melanomas (see **Supplementary Fig 4i**), thus representing putative prospective markers of response. Remarkably, we detected that fewer genes upon loss confer resistance to the nilotinib/trametinib combination compared to trametinib alone (18 vs 29, **Fig 4b**), suggesting that the combination is protective from some molecular mechanisms of trametinib resistance.

In summary, we performed high-throughput drug screenings in melanoma cell lines and found that the combination of nilotinib with trametinib was synergistic in *BRAF*/*NRAS* WT melanoma *in vitro* and *in vivo*. Our results provide a rationale for the clinical development of the nilotinib/trametinib combination.

## Methods

### Cell line

The origin of the melanoma cell lines and the culture medium used to grow them is detailed in **Supplementary Table 1b**. Media was supplemented with 10% Fetal Bovine Serum, Penicillin (100U/ml), Streptomycin (100U/ml) and L-glutamine (292μg/ml) from Gibco. All cell lines were maintained at 37°C and 5% CO_2_ and were tested and found negative for mycoplasma contamination. Cell line identity was confirmed by STR profiling. The nomenclature of PDX lines (and derived samples) include suffixes; indicating the *in vivo* passage number, as described previously^27^.

### Catologue of somatic variants in cell lines

Whole exome sequencing (WES) was performed on 21 melanoma cell lines and the available matched germline (see **Supplementary Table 2b**). DNA libraries were prepared from genomic DNA, exonic regions were captured with the Agilent SureSelect Target Enrichment System, 50 Mb Human All Exon kit or with baits from the Illumina’s TruSeq Exome kit. For C037 whole-genome sequencing (WGS) was performed: libraries were prepared using the standard Illumina library preparation protocol. Paired-end reads of between 70 and 100 bp were generated on the HiSeq 2000 Illumina platform.

WES reads were aligned to the reference genome GRCh37 using the Burrows-Wheeler Aligner software (version 0.7.5a-r406)^38^. MuTect (v1.1.4)^39^, with default parameters, was used to identify somatic point mutations from read alignments by comparing cell lines with matched germlines. For MeWo, Colo-792, CHL-1, M002.X1.CL, D10 lines without a normal germline reference, we used WGS data from C037 germline as reference. The effects of mutations on protein sequences was predicted using the Variant Effect Predictor^40^ and gene models from Ensembl release 75^41^. Each mutation was also annotated with data from the ExAC database (ExAC allele frequency from version 0.3)^40^, and the COSMIC database (mutation ID and number of human tumor samples in COSMIC carrying that mutation, database version 71)^42^. To reduce the number of false somatic mutations in cell lines without matched germline, we removed: 1) single nucleotide variants reported by SAMtools mpileup against the human reference genome (GRCh37) in any of the 16 germline samples in our collection; 2) all the mutations with an ExAC allele frequency >0.5%. We considered all the mutations in splice sites or coding regions and used the Variant Effect Predictor annotation to define missense and loss of function (LOF) mutations according to the following table:

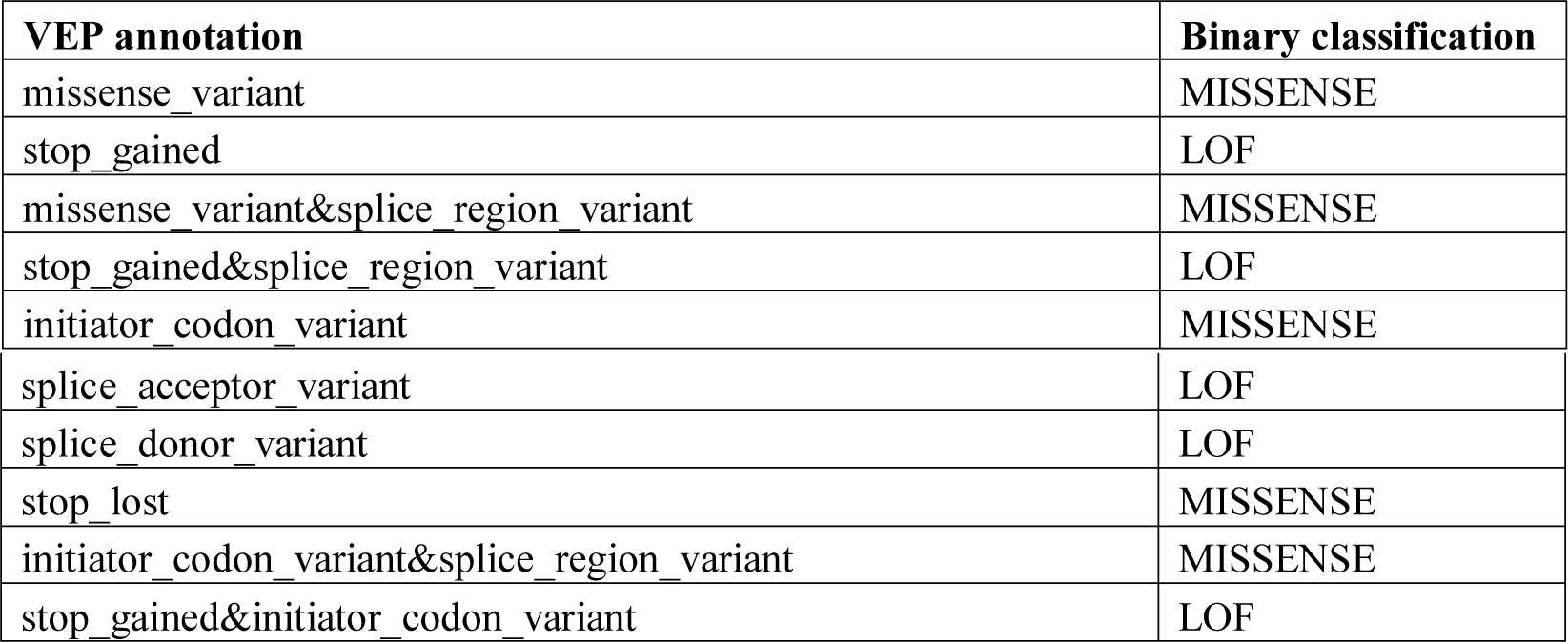

| **VEP annotation** | **Binary classification** |
| --- | --- |
| missense_variant | MISSENSE |
| stop_gained | LOF |
| missense_variant&splice_region_variant | MISSENSE |
| stop_gained&splice_region_variant | LOF |
| initiator_codon_variant | MISSENSE |
| splice_acceptor_variant | LOF |
| splice_donor_variant | LOF |
| stop_lost | MISSENSE |
| initiator_codon_variant&splice_region_variant | MISSENSE |
| stop_gained&initiator_codon_variant | LOF |

These mutations were reported in **Supplementary Table 2a** and are the set of somatic mutations throughout the manuscript.

The *NRAS*^Q61R^ mutation in D38s was identified from the RNA Sequencing data (see “Mutation validation with RNA-seq”) and validated by Sanger sequencing.

To compare data between bait sets used for WES and also WGS, we considered only the mutations within the overlap between Agilent SureSelect Target Enrichment System 50 Mb Human All Exon kit and Illumina’s TruSeq Exome kit. These mutations were used to compile the plots in **Supplementary Figure 1a-d** and **Supplementary Table 2b-c**.

The mutations of the cell lines from the Sanger Institute Cancer Cell Line Panel have been described previously^8^. The *BRAF* and *NRAS* status of the cell lines from the Herlyn’s lab collection have been described previously (https://www.wistar.org/lab/meenhard-herlyn-dvm-dsc/page/melanoma-cell-lines-0).

According to the criteria defined by TCGA for human tumors^4^, we defined as *BRAF*/*NRAS* wild type melanoma lines that do not carry any mutations at these amino acid positions: BRAF^V600^, BRAF^V601^, NRAS^G12^, NRAS^G13^, NRAS^Q61^. None of our cell lines carry any mutation in *HRAS* or *KRAS*. We classified as *NF1* mutant the *BRAF*/*NRAS* wild type melanoma cell lines carrying any non-synonymous mutation in the *NF1* gene (NF1m), and the others as triple wild type (TWT).

### Mutation validation with RNA-seq

RNA-sequencing reads (see “Gene expression analysis by RNA sequencing”) from melanoma cell lines were aligned to the reference genome GRCh37 using the STAR aligner (version 2.5.0)^43^. A 2-pass STAR alignment was performed, and BAM files from replicates were merged. PCR duplicates were flagged using Picard (version 1.135; http://broadinstitute.github.io/picard/) and base quality score recalibration (BQSR) performed using the Genome Analysis Toolkit (GATK; version 3.5)^44^ prior to running the GATK HaplotypeCaller. Sites covered with a minimum of 20 reads were considered for comparison with mutations called from WES. We only considered missense mutations since LOF mutations would likely be associated with unstable mRNA thus leading to an underrepresentation over the wild type allele in the RNA-Seq reads. Most of the cell lines displayed a very high concordance between RNA-seq and exome mutation calls (median 92.5%). Overall, we validated by RNA-seq 15,290 mutations, corresponding to 91.95 of the mutation covered by RNA-seq reads.

### Analysis of copy number variation

Genome wide copy number was determined using the Affymetrix Genome-Wide Human SNP Array 6.0. Data analysis was performed with PICNIC^45^. The copy number information for each gene is presented as (**Supplementary Table 3a**): maximum and minimum copynumber (of any genomic segment containing coding sequence of the gene); zygosity (scored ‘0’, ‘L’ or ‘H’ if any genomic segment is homozygously deleted, has loss of heterozygosity or the whole region is heterozygous, respectively) and disruption status (D, if the gene resides on more than one genomic segment).

### Gene expression analysis by RNA sequencing

The 22 cell lines used for the high-throughput drug screen were grown in biological triplicate (>3 independent passages among replicates, named A-B-C in **Supplementary Table 4**) and collected at 60-80% confluence. Total RNA, including small RNA, was extracted with the microRNAeasy mini kit (Qiagen). We prepared 2μg of RNA for each sample which was spiked with ERCCv92 Mix 1 (Ambion, Thermo Fisher) to measure the dynamic range of detection. Stranded RNA-Sequencing libraries were prepared with the standard Illumina cDNA protocol with a library fragment size between 200 and 300bp. Three multiplexed libraries were prepared each with 22 samples and containing a biological replicate for each cell line. For each sample we obtained ∼ 55 10^6 paired end reads of 100bp on the HiSeq2000 platform.

The RNA sequencing reads were mapped against the human genome (GRCh37d5) using Tophat2^46^ (v2.0.10) and an annotation file containing ENSEMBL v75 with the following parameters (--library-type fr-firststrand -g 1 -G). Subsequently, read pairs were counted using htseq-count from HTSeq^47^, based on the ENSEMBL v75 annotation (Parameters; -m intersection-nonempty -a 10 -i gene_id -s reverse). Using the counts obtained and the average transcript length per gene, we calculated the number of Fragments Per Kilobase per Million reads mapped (FPKM) per gene to assess expression (**Supplementary Table 4a**). RNA-seq data from cell-lines C022 and D35 were too low quality to reliably assess gene expression hence, they were not considered for any further analyses.

### Definition of the differentially expressed genes between sensitive and non-sensitive cell lines

We compared the gene expression of cell lines sensitive to nilotinib/trametinib combination with the gene expression of cell lines non-sensitive to the combination (5 vs 6 cell line with RNASeqd data available, see “Definition of the delta AUC threshold for synergy” for definition). The statistical approach took as input the RNA sequencing reads data from the biological triplicate of each cell line through the following steps. Firstly we considered only those genes that are expressed with FPKM>1 in >2 cell lines of the collection. Then Voom^48^ was used to normalize the read counts for the library size, log-transform the read counts such that the distribution becomes Gaussian-like and estimate precision weights to account for variation in precision between observations^48484848515151515151^. Limma’s^49^ duplicateCorrelation() function was used to incorporate the information from replicates using a mixed modelling framework. Voom was used both before and after duplicateCorrelation(), to normalize the input and to take into account the replicates in the normalization, respectively. Finally, Limma was used to identify differentially expressed genes. We considered as differentially expressed those genes that had a False Discovery Rate (FDR) corrected *P*-value <0.05 and fold change >2 or <0.5.

### Interrogation of the genes differentially expressed in sensitive and non-sensitive cell line in human tumor transcriptome data

The genes differentially expressed between sensitive and non-sensitive cell lines were used to probe the transcriptome of melanoma tumors from TCGA and Leeds Melanoma Cohort (LMC)^50^ using the nearest centroid method^15, 50^. We averaged the 320 genes of the synergy signature within each cell type class (sensitive and non-sensitive), creating a ‘synergy’ and a ‘non-synergy’ centroid vector, each gene having been standardised (mean 0 and variance 1) beforehand. To classify each tumor as synergy-like or non-synergy-like, its standardised expression values (mean 0 and variance 1) were correlated with each centroid. Then the tumors was assigned to the group showing the highest correlation, with at least a difference of0.1 in Spearman correlation coefficients between the 2 groups. A tumor was deemed unclassified if the difference in correlation coefficients was lower than 0.1. A similar approach was used to classify tumors in one of the 4 molecular classes defined by the Jonsson et al. signature (proliferative, pigmentation, high-immune and normal-like^15, 51^). For this analysis, a tumor was deemed classifiable if its Spearman correlation coefficient with one of the 4 classes was greater than 0.1, with the highest correlation coefficient determining the Jonsson’s class to which the sample was allocated.

### Quantitative RT-PCR for *AXL* in PDX samples

RNA from *BRAF*/*NRAS* WT PDX was isolated using Trizol, according to manufacturers’ protocol. cDNA was generated using the Maxima First Strand cDNA Synthesis Kit (Thermo) according to manufacturers’ protocol. Real-time PCR was performed using the following primers:

HPRT-F:5’-CGGCTCCGTTATGGCG-3’;

HPRT-R: 5’-GGTCATAACCTGGTTCATCATCAC-3’;

AXL-F: 5‘-GGTGGCTGTGAAGACGATGA-3’;

AXL-R: 5’-CTCAGATACTCCATGCCACT-3’;

The SYBR-Hi ROX kit (Roche) was used according to manufacturers’ protocol with the Step One Plus Real Time PCR System (Applied Biosystems). *AXL* expression levels were normalized to the *HPRT* housekeeping control.

### MicroRNA expression analysis

Libraries for microRNA sequencing were prepared from the RNA extracted as described above with the Illumina Small RNA library kit. For each sample we obtained ∼ 9 10^6 single end reads of 50bp on the HiSeq2000 platform. The sequencing reads were mapped with Chimera^52^ and Blasted against microRNA precursor sequences obtained from the miRBase version 21 (http://www.mirbase.org/) database. Counts were normalised using DESeq2^53^. To define up or downregulated microRNAs, we compared each cell line vs all the other cell lines within the collection and calculated statistical significance of the difference with Voom^48^. We considered significant those microRNAs with a Voom t-statistic value >10 or <-10, and with an absolute value of the log_2_ fold change >v2. If more than 50 microRNAs resulted, we selected only the top 50 microRNAs (by T value ranking). If fewer than 5 microRNAs resulted, we selected the top 5 regardless of the threshold criteria.

### Definition of *BRAF/NRAS* WT melanoma drivers

To identify cancer drivers specific for *BRAF/NRAS* WT disease, we analysed the mutation calls from 74 *BRAF/NRAS* WT melanomas (i.e. NF1 mutant plus triple wild type) from the TCGA collection^4^ using the IntOGEn pipeline^10^ with the 3 algorithms Mutsig, OncodriveClust and OncodriveFM. We considered as mutation drivers those genes that have a significant signal (Q-value<0.05) with one of the 3 algorithms and that are known drivers in other tumor types. Given the difficulty to identify recurrently amplified and deleted genes from a small collection of samples such as the *BRAF*/*NRAS* WT melanomas, we considered as CNV melanoma drivers the genes that map within chromosomal regions previously defined as significantly amplified or deleted in the TCGA melanoma collection (n= 333)^4^.

### High throughput drug screening

We tested a library of 60 drugs targeting the main pathways deregulated in cancers. The range of the drug concentrations was defined according to the activity of each compound against a large panel of cell lines^8, 54^ (see **Supplementary Table 8** for drug description, supplier and concentrations used). The 3 anchor drugs temozolomide, nilotinib and roscovitine were used at 2 different concentrations.

Cell lines were seeded in 384-well microplates at low confluency in culture medium. The optimal cell number for each cell line was determined to ensure that each was in growth phase at the end of the assay (∼85% confluency). After overnight incubation cells were treated with 5 concentrations of each compound (4-fold dilution series, covering a 256-fold drug concentration range), using liquid handling robotics (Beckman Coulter), and then returned to the incubator for 6 days. At day 6, cells were fixed in 4% formaldehyde for 30 minutes, then stained with 1μM of Syto60 red fluorescent nucleic acid stain (Molecular Probes, Thermo Fisher) for 1 hour. Quantitation of fluorescent signal intensity was performed using a fluorescent plate reader at excitation and emission wavelengths of 630/695nm. All screening plates were subjected to stringent quality control measures and to assess the quality of our screening a Z-factor score comparing negative and positive control wells was calculated across all screening plates.

### Analysis of high-throughput viability data

We derived the Area Under the Curve (AUC) parameter from the cell line viability data normalized for vehicle treated control. Empirical values above 1 or below 0 were capped to values of 1 or 0, respectively. The AUC was computed using a trapezoid integration below the 5 measured viability values of the dose-response curve. We calculated the AUC for the library drug, anchor drug and drug combination.

We derived the delta AUC value to measure drug synergy. We used the Bliss independence model^11^ to compute the expected viability of the cell line when exposed to the drug pair (as arithmetic product of the viability measured with the library drug alone and the viability measured with the anchor drug alone). This defined the expected dose-response curve on the 5 measured concentrations of the library drug used in combination with the anchor drug. Throughout the manuscript we called the predicted AUC of the combination defined by the Bliss model as predicted additivity. The delta AUC is defined as the difference between theAUC below the predicted additivity dose-response curve and the AUC below the experimentally observed dose-response curve.

### Low throughput viability assays

We performed low throughput viability assays to validate the results obtained with the high-throughput drug screening. Each experimental point was performed in technical triplicate. We used the same anchor drug concentration and the same 256 fold range of library drug concentrations of the high-throughput drug screening, but with 9 points with a 2-fold dilution series for the library drugs.

The cell lines were seeded in 96 wells-microplates at a non-saturating density (confluency 60-90% after the 6 days in vehicle treated control, see **Supplementary Table 10d**) were drugged them the day after. At 6 days the cells were fixed in 4% formaldehyde and staining was performed with 1μM Syto 60 red fluorescent nucleic acid stain (Molecular Probes,Thermo Fisher). The MW96 plates were read using a Biomek FXc Liquid HandlingAutomation Workstation (Beckman Coulter). The analysis of the AUCs and delta AUCs was performed as described above for the high-throughput drug screening.

### Clonogenic assays

We seeded in each well of a 6 well microplates the same number of cells used for the low throughput viability assays in 2ml of media. 24h after seeding we added 2ml of media containing the dilution of the drug(s). After 10-15 days of drug treatment, when clones became evident, the cells were fixed with methanol for 1h, then stained for 30 seconds with 0.5% crystal violet (Sigma-Aldrich) dissolved in 25% methanol and washed twice in water. The assays were performed for representative cell lines that grew efficiently at the required low density.

### Definition of the delta AUC threshold for synergy

In order to triage the drug combinations for validation, firstly we selected the five drug combinations with the top average delta AUC in the high-throughput screening that met the criteria of having a delta AUC synergy score >0.2 in three or more cell lines. Priority was given to the highest dose of anchor drug as this resulted in increased activity of the combination. The threshold of delta AUC>0.2 was selected as it corresponds to the top 1.5% delta AUC of all the screened drug combinations. In an effort to extend the pool of validated drug combinations, we also tested the three combinations with delta AUC>0.2 in two cell lines only, displaying the highest activity (AUC<0.4). None of those three combinations was successfully validated in any of the cell lines, suggesting that decreasing the threshold is unlikely to identify reliable hits.

Biological replication of the low throughput assay (n=8) in C077 cell lines showed high reproducibility, with a standard deviation of the delta AUC of 0.0475267. We therefore considered as synergistic those drug combinations that displayed a delta AUC>0.1 in the low throughput assays, a value that is above 2 fold the standard deviation of the assay. The agreement among the biological replicates of the so-defined synergy confirmed the reliability of the delta AUC threshold (**Supplementary Table 11**).

To flag cell lines where the drug combination achieved a high killing activity (**Fig. 2d**), we used an AUC combination threshold below 0.4, a value that in the high-throughput screens represents the bottom 5% values of AUC combination (i.e. the combination with the top 5% activity).

For 21 melanoma cell lines we tested nilotinib combined with 2 MEK inhibitors (trametinib and PD-0325901). We classified as ‘sensitive’ those cell lines displaying a delta AUC>0.1 for both nilotinib plus trametinib and nilotinib plus PD-0325901. We classified as ‘non-sensitive’ those cell lines that displayed a delta AUC<0.1 for both nilotinib plus trametinib and nilotinib plus PD-0325901. The cell lines which displayed delta AUC>0.1 for one of the MEK inhibitors combined with nilotinib and a delta AUC<0.1 for the other MEK inhibitor were classified as “intermediate”. The cell lines that displayed an AUC<0.3 for the anchor drug alone or the library drug alone were classified as synergy not detectable (“ND”), since the high activity of a single drug alone hampered the reliable detection of synergy (see **Supplementary Table 10b-c**).

For the definition of synergy and for the association with mutation, CNV, gene and microRNA expression data, we used the data from the low throughput validation described in **Supplementary Table 10b**.

The observed synergies were tested in biological replicates (range of biological replicates 2-8, **Supplementary Table 11 a-b**); at least 2 biological replicates were performed by 2 different operators.

### Definition of the cell lesions used for the association with the drug sensitivity data

We collected a list of high confidence cancer driver lesions defined by previous studies^55, 56, 57, 58, 59, 60^ (**Supplementary Table 7d**), following an approach previously successful in the identification of drug sensitivity markers^8^. To generate the list of driver lesions in melanoma driver genes, we collated all the somatic LOF mutations in the 24 *BRAF*/*NRAS* WT melanoma drivers (see “Definition of *BRAF/NRAS* WT melanoma drivers” and**Supplementary Table 2**) and selected only the somatic missense mutations that matched the previously defined list of high confidence cancer driver lesions (**Supplementary Table 7d**). To extend the list of considered lesions, we compiled a list of all lesion in melanoma driver genes considering all the LOF and all the missense somatic mutations in any position within the 24 *BRAF*/*NRAS* WT melanoma drivers (**Supplementary Table 7b**). We then defined the list of copy number alterations in our cell lines by considering only the 39 genes in region significantly amplified/deleted in melanoma and the 24 *BRAF*/*NRAS* WT melanoma drivers genes (**Supplementary Table 7b-c**). We defined each of those genes as amplified (AMP) if the gene is in a segment with >5 copies or deleted (DEL) if the gene is in a segment with <2 copies according to the SNP6 array data analysis with PICNIC algorithm (**Supplementary Table 3a**).We defined the list of up and downregulated genes for these 24 *BRAF*/*NRAS* WT drivers and the 39 genes in region significantly amplified/deleted in melanoma by 1) averaging the biological triplicate per cell line; 2) removing the genes that have FPKM>1 in less than 3 cell lines (poorly expressed genes); 3) dividing the cell line specific FPKM expression value of each gene for the median of expression of that gene in the whole collection; 4) defining as upregulated (UP) those genes with a fold change over the median>4 and as downregulated those genes with a fold change over the median <0.25. We defined the up/downregulated microRNAs as described above (see “MicroRNA expression analysis”). We then summarized by gene the two versions (driver lesions in driver genes and all lesions in driver genes) of cell lesions and analysed their association with drug synergy data (**Supplementary Table 12 a-b**). For both sets we considered the different types of lesions combined together, as a single input, or grouped according to putative functional impact, as detailed in **Supplementary Table 13d**. For the identification of the statistical association with the drug synergy score, identical alteration profiles were merged. Only genes with a lesion in >2 cell lines were considered for the analysis. For each cell lesion defined as detailed above, we compared the delta AUC score between the cell lines with or without the lesion using a t-test. The p-values obtained were corrected for multiple testing with the Benjamini-Hochberg method. Statistical analyses were performed with R/Bioconductor^61^.

### Western blot analysis

To measure the level of p-ERK upon drug treatment, cells were seeded at twice the density of that used for low throughput viability assays (**Supplementary Table 10d**). They were drugged the day after, and proteins were collected 6h later using NP40 lysis buffer (Thermo Fisher Scientific) containing Protease/Phosphatase Inhibitor Cocktail (Cell signalling). Protein lysates were quantified with Pierce BCA Protein Assay kit (Thermo Fisher Scientific).

Proteins were denatured by adding 25% of NuPAGE LDS Sample Buffer (Thermo Fisher Scientific) and 5% of dithiothreitol 1M (Sigma) and incubating 15 minutes at 75°C. 5-10μg of protein were loaded on NuPAGE™ Novex™ 4-12% Bis-Tris Protein Gels (Thermo Fisher Scientific) and electrophoresis was performed at 120V in NuPAGE® MOPS SDS Running Buffer with NuPage Antioxidant (both from Thermo Fisher Scientific) in a Xcell Surelock electrophoresis cell. Proteins were transferred to Amersham Hybond N+ nylon membrane (GE Healthcare) by overnight blotting at 4°C at 10V in XCell II blot machine (Lifetech) in NuPAGE Transfer Buffer with NuPage Antioxidant (both from Thermo Fisher Scientific). Membrane blocking was performed in 5% non-fat milk (Cell Signalling) or 5% BSA (Acros Organics) dissolved in Tris buffered saline with 0.25% of Tween 20 (TBS-Tween, Sigma-Aldrich, see table below). Antibodies usage is described in the table below

| target protein | catalogue number | company | dilution | blocking | species |
| --- | --- | --- | --- | --- | --- |
| AXL for cell lines | #8661 | Cell Signalling | 1:1000 | BSA | rabbit |
| MITF | ab12039 | abcam | 1:1000 | BSA | mouse |
| HSP-90 | #4875 | Cell Signalling | 1:2000 | BSA | rabbit |
| total ERK | #9102 | Cell Signalling | 1:2000 | BSA | rabbit |
| P-ERK | #4370 | Cell Signalling | 1:6000 | BSA | rabbit |
| beta tubulin | #5346 | Cell Signalling | 1:3000 | BSA | rabbit |
| vinculin | V9131 | Sigma | 1:1000 | milk | mouse |
| AXL for PDX | sc-20741 | Santa Cruz Biotechnology | 1:1000 | milk | rabbit |

Membranes were incubated overnight at 4°C with the primary antibody. After washing with TBS-Tween, incubation with secondary antibody (anti-rabbit or anti-mouse IgG HRP-linked (1:6000 and 1:3000, #7074 and #7076, respectively, from Cell Signalling) was performed at RT for 1h. The membrane was washed with TBS-Tween and the signal detected with Amersham ECL Select Western blotting detection reagent (GE Healthcare) using Image Quant Las4000.

Immunoblotting for PDX samples was performed following the protocol previously described^62^.

The signal on the western blots images were quantified using ImageJ (Rasband, W.S., ImageJ, U. S. National Institutes of Health, Bethesda, Maryland, USA, http://imagej.nih.gov/ij/, 1997-2016.)”. Each gel lane was outlined and the densitometry plotted. The peak of interest was then defined and quantified. Each value obtained was normalised to a loading control. Each experiment for the detection of P-ERK levels was performed in biological triplicate. The analysis of the images was performed by an operator blinded to the sample identifiers.

### AXL overexpression and knockdown

We used two lentiviral vectors expressing two short hairpin RNAs (shRNA1 and shRNA3) targeting the AXL transcript (from theTRC shRNA library, Dharmacon). A luciferase targeting shRNA sequence in the same lentiviral backbone was used as a negative control. The lentiviral vector used to overexpress AXL was generated by cloning the human AXL ORF into the pCDH backbone. The empty pCDH backbone was used as negative control. Lentiviral vectors were produced as previously described^63^. The melanoma cell lines were infected with different volumes of lentiviral vectors as indicated throughout the manuscript to account for the different viral titers and the different transducibility of each cell lines. Three days after infections cells were selected with puromycin 2 μg/ml for 1-2 weeks. Proteins were collected for Western blot analysis and low throughput viability assays were performed as detailed above.

### CRISPR/Cas9 screening

We performed CRISPR/Cas9 screens to identify mechanisms of resistance to the nilotinib/trametinib combination (see outline in **Supplementary Fig. 4a**). We used a previously described genome-wide library of synthetic guide RNAs (sgRNA) targeting the human genome (library V1, containing 90,709 sgRNA targeting 18,010 human genes^22^) for the CHL-1 cell line, and an updated version of the same library (library V1.1) for C077 and MeWo. The generation of Cas9 expressing cell lines, measurement of Cas9 activity and virus titration was performed as described previously^22^. For each cell line we infected 60 10^6 cells in duplicate at multiplicity of infection 0.3 (200X library representation in each replicate). Selection with puromycin 2 μg/ml was carried out for 4 days and 2 weeks after infection 60 10^6 cells per replicate were seeded in drug regimens: trametinib (100nM for CHL-1; 12nM for C077 and MeWo), nilotinib (2μM for all the lines) plus trametinib (100nM for CHL-1; 12nM for C077 and MeWo) and matched DMSO as a control. Given the limited activity of nilotinib alone (**Supplementary Table 10b**) which would have resulted in the absence of selective pressure, we did not screen using nilotinib alone. An aliquot of 60 10^6 was collected before the start of drug selection as a reference population (PRE population, whose resulting sgRNA counts were used to calculate the ROC curve, see below). Cells were split when confluent maintaining >60 10^6 cells per population. After 18 days of drug selection,>60 10^6 cells were collected from each cell line population. DNA was extracted with the Qiagen Blood & Cell Culture DNA Maxi Kit (Qiagen). Amplification of sgRNA and sequencing was performed as described^22^, starting with 72 μg of template DNA and obtaining∼ 50 10^6 sequencing reads per sample. To measure reproducibility, we compared the sgRNA normalized counts between the replicates of infection before the drug administration (PRE population) (R = 0.74-0.93). As a quality control we estimated the ability of each CRISPR/Cas9 screen to discriminate between genes belonging to known sets of essential and non-essential genes^64^, *E* and *N* respectively. To do this we aggregated sgRNA depletion p-values through MAGeCK, yielding a gene level summary of essentiality. The genes were then sorted according to their gene-level depletion p-values. At each gene rank position in the sorted list we compiled the true positive rate (fraction of genes belonging to E) and false positive rate (fraction of genes belonging to N), and created a receiver operating characteristic (ROC) curve plotting Sensitivity vs. (1-Specificity) at each rank. The area under the ROC curve for the 3 cell lines was >0.9, indicative of a successful screen. Finally, we found that none of the significantly enriched genes with FDR<0.1 in both replicates of each cell lines (i.e. what we defined as hits) were poorly expressed genes (FPKM<1)), indicating the high specificity of the screen.Each replicate of infection and selection (with trametinib or drug combination) was compared with the sister DMSO treated control. MAGeCK^65^ was used to identify genes whose sgRNA targeting pool was significantly enriched or depleted compared to the control. We considered as significant hits those genes with FDR corrected p-value <0.1 in both the replicates per cell line.

### Enrichment and protein-protein network analysis

The enrichment analysis was performed by MsigDB (http://software.broadinstitute.org/gsea/msigdb/annotate.jsp) considering Canonical pathways and Hallmark gene sets and the top 100 pathways with FDR<0.05. The database was accessed in December 2016. The Network of protein-protein interactions were defined using STRING^23^ (http://string-db.org). We interrogated multiple proteins from Homo Sapiens gene symbols and to obtain a protein network image. We also used the output of the Gene Ontology enrichment analysis from STRING and integrated with the MsigDB output described above to display the top enriched protein complex or pathways in **Fig 4c**. The STRING database was accessed in January 2017.

### Interrogation of the status of the drug resistance genes identified from CRISPR/Cas9 screening in melanoma

We interrogated the cBioportal database (http://www.cbioportal.org/) in March 2017 to investigate the status of the 18 nilotinib/trametinib resistance genes found as significant hits of the CRISPR/Cas9 screening in ≥2 cell lines. Four datasets of skin melanoma were available: Broad Cell 2012 (121 samples); Broad/Dana Faber, Nature 2012 (25 samples); TCGA, provisional (287 samples); Yale, Nat Genet 2012 (91 samples). CCDC101 genesymbol was present in the database with the alternative symbol SGF29. We reported the frequency of samples with an alteration in one of those genes in the four datasets, and provided detailed mutation type for the 2 larger dataset (TCGA and Yale). Only the TCGA dataset included copy number variation data.

### Interrogation of previously published screen in melanoma drug resistance

We interrogated the GenomeCRISPR website (http://genomecrispr.dkfz.de/) for Cas9 drug resistance screens in melanoma in (January 2017. We found 2 studies that performed genome-wide Cas9 screenings in A375 (*BRAF*^V600E^-mutant) with vermurafenib (BRAF inhibitor) and another screen that used selumetinib (MEK inhibitor) ^24, 25^. We compiled a list of vemurafenib resistance genes considering the hits identified by Shalem et al^24^ that were found as in the top 100 ranking genes by RIGER score in both the replicates; to this list we added 2 new hits found by Li et al.^65^ following a re-analysis of the same data with MAGeCK. We then included the list of vemurafenib resistance genes identified by Doench et al^25^ with the GeckoLV2 library with a FDR corrected P-value from the STARS algorithm<0.1.

We compiled a list of selumetinib resistance genes with the gene genes identified by Doench et al^25^ with the GeckoLV2 library that have a FDR corrected P-value from STARS algorithm<0.1.

### Mass spectrometry for proteomics and phosphoproteomics analyses

Proteomics and phosphoproteomics analysis was performed as previously described with minor modifications^66^. The cell pellets were dissolved in 0.1 M triethylammonium bicarbonate (TEAB), 0.1% SDS, 10% isopropanol with pulsed probe sonication (EpiShear™, power 40%) on ice for 20 sec and direct boiling at 95 °C. Protein concentration was measured with Quick Start Bradford Protein Assay (Bio-Rad). Cysteine disulfide bonds were reduced with tris-2-carboxymethyl phosphine (TCEP) and cysteine residues were blocked with Iodoacetamide (IAA). Trypsin (Pierce, MS grade) was added at mass ratio 1:30 for overnight digestion. The resultant peptides were labelled with the TMT 10-plex reagents (Thermo Scientific) according to manufacturer’s instructions. Samples were combined and the mixture was dried with speedvac concentrator and stored at -20 °C. High pH Reverse Phase (RP) peptide fractionation was performed with the Waters, XBridge C18 column (2.1 x 150 mm,3.5 μm, 120 Å) on a Dionex Ultimate 3000 HPLC system over a 35 min gradient. Fractions were collected every 30 sec and were dried with SpeedVac concentrator. The peptide fractions were reconstituted in 10 uL of 20% isopropanol, 0.5% formic acid binding solution and were loaded on 10 uL of phosphopeptide enrichment IMAC resin (PHOS-Select™ Iron Affinity Gel) already conditioned with binding solution. The resin was washed three times with 40 uL of binding solution and centrifugation at 300 g after 2 h of binding and the flow-through solutions were collected. Phosphopeptides were eluted three times with 70 uL of 40% acetonitrile, 400 mM ammonium hydroxide solution. Both the eluents and flow-through solutions were dried in a speedvac and stored at -20 °C until the phosphoproteomic and proteomic LC-MS analysis respectively. LC-MS analysis was performed on the Dionex Ultimate 3000 UHPLC system coupled with the Orbitrap Fusion Tribrid Mass Spectrometer(Thermo Scientific). The peptide fractions were subjected to separation on the Acclaim PepMap RSLC (75 μm × 50 cm, 2 μm, 100 Å) C18 capillary column over a 95 min gradient.Precursors were selected with mass resolution of 120k in the top speed mode and were isolated for CID fragmentation with collision energy 35%. MS3 quantification spectra were acquired with Synchronous Precursor Selection (SPS) and 50k resolution. Phosphopeptide samples were analyzed with a top15 HCD method at the MS2 level. The acquired mass spectra were submitted to SequestHT search in Proteome Discoverer 2.1 for protein identification and quantification. The precursor mass tolerance was set at 20 ppm and the fragment ion mass tolerance was set at 0.5 Da for the CID and at 0.02 Da for the HCD spectra used for the phosphopeptide analysis. TMT6plex at N-termimus, K and Carbamidomethyl at C were defined as static modifications. Dynamic modifications included oxidation of M and Deamidation of N, Q. Search for phospho-S,T,Y was included only for the IMAC data. Peptide confidence was estimated with the Percolator node. Peptide FDR was set at 0.01 and validation was based on q-value and decoy database search. All spectra were searched against a UniProt fasta file containing 20,165 reviewed human entries. The Reporter Ion Quantifier node included a TMT-10plex Quantification Method with integration window tolerance 15 ppm at the MS3 level or at the MS2 level for the IMAC data. Only peptides uniquely belonging to protein groups were used for quantification. Phosphopeptides with signal/noise ratio<5 and carrying oxidation or deamidation were not considered for the analyses. The FDR corrected *P*-value presented in Supplementary Table 17 were calculated with the function software ‘Perseus’ using the ‘Significance’ function, considering the phosphopeptides that represents significant outliers in the variations vs the DMSO-treated controls and among all the phopshopeptides within the sample. We considered the list of nilotinib targets as published by Davis et al^18^.

### *In vivo* animal experiments

Animal experiments were approved by the animal experimental committee of the Netherlands Cancer institute and performed according to Dutch law in the Netherlands. PDX were generated as described^28^. We injected 70,000 (M003.X2) – 300,000 (MeWo) cells in both flanks of NOD.Cg-*Prkdc*^scid^ *Il2rg*^tm1Wjl^/SzJ (NSG) mice. After the tumors reached ∼50-90mm^3^ volume, mice were randomized in 4 treatment groups (n=5 mice/group, n=10 tumors/group) that were administered by oral gavage: a) vehicle (0.5% hydroxypropylmethyl cellulose (HPMC, Sigma) aqueous solution containing 0.05% Tween 80); b) nilotinib 75mg/kg/day (administered 37.5mg/kg twice daily), c) trametinib 0.1-0.3 mg/kg/day for MeWo and M003.X2, respectively; d) nilotinib plus trametinib combination. Mouse weight was monitored weekly; tumor size was measured by caliper 3 times per week. Mice were euthanized either when the tumor volume reached 1000mm^3^ or when the weight loss of the mice was more than 30%, or at the experimental endpoint.

### Statistical analyses

Graphs and statistics were generated using the GraphPad Prism software. The statistical test applied is indicated in the respective Figure legends. Briefly, when two groups were analysed, the *P-*value was calculated by unpaired Student’s t-test; when three or more groups were analysed, the *P-*value was calculated by one way Anova and Tukey’s multiple comparisons test. For the tumor growth curve *in vivo*, the group treated with the drug combination was compared to each of the other 3 groups by unpaired Student’s t-test considering the tumor volumes measured at the last experimental time point when all 4 cohorts were viable. A two tailed Fisher’s exact test was used to compare the number of tumors classified as synergy like and non-synergy like between one of the 4 Jonsson’s expression classes and the remaining 3 classes (**Supplementary Fig 3c-d**), as well as to assess the association between AXL expression and nilotinib/trametinib synergy (**Supplementary Table 15c**).

### Data Availability

The WES data (and WGS data for C037), the SNP6 array raw data (for CNV) and the RNA sequencing data for the first collection of cell lines are deposited in ArrayExpress with Accession number (E-ERAD-293) and ENA, with accession number (EGAS00001000815). The small RNA sequencing data are deposited in ArrayExpress with Accession Number (E-ERAD-294) and ENA with Accession Number (EGAS00001000816).

For the interrogation of the gene expression pattern associated to synergy in the transcriptome of human tumors, the data from the extended TCGA cohort of melanoma (n=474) were downloaded from TCGA database in early 2016 as RSEM values; the RSEM values that we used for our analysis can be downloaded from the following link: ftp://ftp.sanger.ac.uk/pub/users/vvi/tcga_rnaseq_v2_level3_skcm/tcga_sckm_rnaseqv2.tsv.gz. The data from the Leeds Melanoma Cohort (LMC) have been published previously^50^.

The mass spectrometry proteomics data have been deposited to the ProteomeXchange Consortium via the PRIDE partner repository with the dataset identifier PXD007649. Temporary reviewer username and password p0R7pMF5.

## ACKNOWLEDGMENTS

We would like to thank all the members of David Adams’, Mathew Garnett’s and Ultan McDermott’s groups for the helpful scientific discussions. This work was supported by: the Cancer Research UK, ERC Combat Cancer Grant, IEF Marie-Curie Fellowship and the Wellcome Trust.

## AUTHOR CONTRIBUTIONS

Mar.Ran. and D.A. conceived the study and wrote the manuscript. Mar.Ran. designed and performed most of the experiments and analyzed data. K.K. and Ai.Sh. and D.P. performed *in vivo* experiments. M.M, O.K., N.A., V.I., K.W., M.D.C., J.N., Mam.Ras., D.T., F.I., S.vD.,G.B., C.A., A.E., J.N., L.W. performed bioinformatics analysis on the data generated and on previously published datasets. An.Sp., G.T., N.T., M.S., E.S., C.W., S.C., T.R., J.C. performed experiments. V.G., U.M. and F.B. generated reagents for the CRISPR/Cas9 screening. K.Y. generated the library for the Cas9 screening. K.D., A.P., N.H. generated melanoma cell lines and mutation data. M.G. performed the high-throughput drug screening. All the authors revised the manuscript.

